# A high-quality, chromosome-scale genome assembly of the shade-tolerant wild rice, *Oryza granulata*

**DOI:** 10.64898/2026.04.28.721348

**Authors:** Fen Zhang, Yue-hong Yang, Wei Li, Cong Shi, Xun-ge Zhu, Li-zhi Gao

## Abstract

*Oryza granulata* Nees et Arn. ex Watt, a diploid wild rice (GG genome), possesses exceptional shade tolerance and is a key genetic resource for rice improvement. However, previous genome assemblies lacked continuity and completeness. Here we present a chromosome-scale reference genome of *O. granulata* using PacBio SMRT (113×), Hi-C (95×), and Illumina sequencing. The final assembly is ∼764.24 Mb, with a scaffold N50 of ∼59.32 Mb, and ∼96.47% of the sequence anchored to 12 chromosomes. BUSCO completeness is ∼98.6%. We annotated ∼42,064 protein-coding genes, of which ∼95.39% were functionally annotated, along with ∼73.46% repetitive elements. The genome assembly and raw sequencing data are available at NGDC (PRJCA061980), NGDC GSA (CRA041662), and NGDC GWH (GWHISVE00000000.1). This high-quality genome will serve as a fundamental resource for evolutionary genomics, conservation biology, and breeding of shade-tolerant rice cultivars.

## Background & Summary

Rice (*Oryza sativa* L.) is one of the most important staple crops globally, serving as a primary food source for more than half of the world’s population^1^. With the projected increase in global population, rice production must double by 2050 to meet future food demands^2^. However, current production bottlenecks, including limited genetic diversity in cultivated varieties, have hindered yield improvements^3^. Furthermore, as one of the most water-consuming cereals, rice cultivation faces mounting pressure from global water scarcity^4^. To overcome these challenges, it is essential to explore the genetic potential hidden in wild rice species, which serve as valuable reservoirs of agronomically important traits such as disease resistance, abiotic stress tolerance, and environmental adaptability^5^.

The genus *Oryza* comprises a rich diversity of species, providing a vast genetic reservoir. Of the 27 recognized species, 25 are wild while 2 represent the independently domesticated cultivated species^1,6^. Based on morphological, cytological, and molecular features, members within the genus *Oryza* are sub-classified into 11 genome types, including six diploids (AA, BB, CC, EE, FF, and GG), and five tetraploids (BBCC, CCDD, HHJJ, HHKK, and KKLL)^7^. Among them, *O. granulata*, a diploid species with the GG genome type, occupies a basal phylogenetic position within the genus and is considered one of the oldest extant wild rice lineages, estimated to have first diverged from other members of the genus^8^. It is naturally distributed in shaded upland habitats in South and Southeast Asia, including China, India, Myanmar, and Thailand^6^. *O. meyeriana*, which is considered to be a closely related species to *O. granulata*^9^, also possesses a GG genome and exhibits similar ecological and genomic characteristics. Together, they form the *O*.

*meyeriana* complex within the lower cladogram of the *Oryza* phylogeny^8^. Unlike most *Oryza* species, which prefer aquatic or moist environments, *O. granulata* and *O. meyeriana* thrive in well-drained soils under forest canopies, exhibiting remarkable shade tolerance^8,10^. This unique adaptation to low-light conditions makes *O. granulata* an exceptional genetic resource for breeding shade-tolerant rice varieties, a trait of increasing importance for sustainable agricultural systems such as agroforestry and intercropping, which can enhance land-use efficiency and resilience to climate change^10,11^.

Despite its agronomic potential, genetic studies and breeding applications of *O. granulata* and *O. meyeriana* have been limited due to the lack of high-quality reference genomes. *O. granulata* possesses the second largest genome among diploid *Oryza* species, estimated at ∼882 Mb^12^, nearly twice the size of the cultivated rice genome (∼389 Mb)^11,13^. A recent chromosome-scale assembly of *O. meyeriana* reported a genome size of ∼761.1 Mb with a contig N50 of 30.46 Mb and 41,989 annotated genes, of which 73.04% were repetitive sequences^9^. Previous efforts to sequence the *O. granulata* genome have yielded valuable insights. For instance, Wu et al. (2018) produced a ∼777 Mb assembly using PacBio and fosmid sequencing, revealing the role of LTR bursts in genome expansion and the absence of tandem repeats in centromeric regions, and identified positively selected genes related to photosynthesis and energy metabolism that may underpin its shade tolerance^10^. Similarly, Shi et al. (2020) published a 736.66 Mb draft genome anchored to 12 pseudo-chromosomes using Hi-C data, providing the first chromosome-scale assembly for the species^14^. However, these existing assemblies suffer from limitations that hinder their utility in functional genomics and comparative studies. The assembly by Wu et al. (2018) was highly fragmented, with a contig N50 of only 262 kb, and lacked chromosome-level scaffolding^10^. Although Shi et al. (2020) improved continuity and incorporated Hi-C data, their assembly was still incomplete, with a scaffold N50 of 916.3 kb and ∼13.6 Mb of sequences remaining unmapped^14^. Both assemblies exhibited gaps in repetitive regions and centromeric sequences, limiting their accuracy in structural variant analysis and TE annotation. These shortcomings underscore the need for a more contiguous, complete, and chromosome-resolved genome assembly to fully exploit the genetic and evolutionary insights offered by *O. granulata*, particularly for identifying and harnessing the key genes governing its shade tolerance and other adaptive traits.

In the last decade, great progress has been made in the comparative genomics of cultivated rice and its wild relatives at the chromosome scale^9,15–19^, highlighting the necessity of a high-quality *O. granulata* genome for comprehensive genus-wide analyses. From an evolutionary perspective, it facilitates the comparative genomics across the *Oryza* genus, enabling the reconstruction of phylogenetic relationships and the identification of lineage-specific genetic innovations^9^. For example, the expansion of gene families related to photosynthesis and energy metabolism in *O*.

*granulata* may underlie its adaptation to low-light environments^10^. Our studies on the evolution of *Oryza* chloroplast genomes have revealed that natural selection acting on photosynthesis-related genes has played a crucial role in promoting adaptation to diverse ecological habitats, including the shaded upland environments characteristic of *O. granulata*^8^. This underscores the potential for leveraging both nuclear and organellar genomic resources to fully understand and utilize the shade adaptation mechanisms of this wild species. Such insights are critical for developing next-generation rice cultivars capable of thriving in shaded environments, thereby supporting the expansion of agroforestry systems and improving crop resilience under the canopy cover increasingly prevalent in changing climates. From a conservation standpoint, *O. granulata* is classified as an endangered species due to habitat loss, rapid deforestation, and human disturbance^20,21^. Previous population genetic studies revealed that this species possesses fairly low levels of genetic diversity within populations but high genetic differentiation among geographically isolated populations^22,23^. Efficient conservation strategies are urgently required to sustain the quality and integrity of its gene pool. The considerable genomic diversity detected through pan-genome analysis demonstrates that *de novo* assembly of genomes from genetically distinct populations helps reveal the origin and evolutionary forces shaping population structure^24^. Thus, a high-quality chromosome-scale genome provides a foundational resource for assessing genomic diversity, identifying adaptive loci, and guiding conservation strategies. It supports the precise identification of candidate genes for shade-tolerant trait introgression into cultivated rice, leveraging wild genetic diversity for breeding. The ability to precisely identify and transfer shade tolerance and drought resistance genes from *O. granulata* into elite cultivars represents a promising strategy for enhancing the sustainability and productivity of rice farming systems worldwide.

Here, we present a high-quality, chromosome-scale genome assembly of *O. granulata* using a combination of long-read PacBio sequencing and Hi-C chromatin interaction data (∼113× PacBio SMART reads and ∼95× Hi-C reads) (**Fig. 1A; Table S1**). *K-mer* analysis showed that the genome size of *O. granulata* was estimated to be ∼798.12 Mb, with a relatively low genomic heterozygosity of ∼0.505% (**Fig. 1B**). Our assembly achieves a scaffold N50 of ∼59.32 Mb (**Table 1**), resulting in the final assembled genome of ∼764.24 Mb. Based on the karyotype of the species (2n=24), approximately ∼96.47% of the contig reads were anchored to 12 pseudochromosomes (**Fig. 1C**; **Fig. 1D**; **Table 1**; **Table S3**), significantly improving continuity and completeness compared to previous versions^10,14^. Benchmarking Universal Single-Copy Orthologs (BUSCO) analysis showed a BUSCO score of 98.6%, further demonstrated the high-quality of the assembled genome (**Fig. 2**; **Table 1**; **Table S4**). The annotation of the *O. granulata* genome showed a repeat sequence proportion of ∼73.46% (∼561 Mb) (**Table S5**). We also provide comprehensive annotations of protein-coding genes, non-coding RNAs, and repetitive elements, including LTR retrotransposons that have shaped the species’ genome architecture. A total of 42,064 protein-coding genes were predicted (**Table 1**), of which ∼95.39% were functionally annotated (**Fig. 3**; **Table S6**). In addition, 651 miRNAs, 937 rRNAs, 994 tRNAs, and 608 snRNAs in the *O. granulata* genome were annotated (**Table 1**).

**Table 1.** Statistic of assembly and annotation of the *O. granulata* genome.

| **Assembly** |  |
| --- | --- |
| Genome size (bp) | 764,240,925 |
| Number of scaffolds | 235 |
| GC content of the genome (%) | 46.28% |
| Contig N50 (bp) | 59,318,324 |
| Scaffold N50 (Mb) | 59.32 |
| Scaffold L50 | 6 |
| Scaffold N90 (Mb) | 50.61 |
| Scaffold L90 | 11 |
| Maximum scaffold length (Mb) | 80.2 |
| Chromosome numbers | 12 |
| Complete ratio of BUSCO (%) | 98.6 |
| **Annotation** |  |
| Protein-coding genes | 42,064 |
| miRNA | 651 |
| tRNA | 994 |
| rRNA | 937 |
| snRNA | 608 |

**Fig 1.**
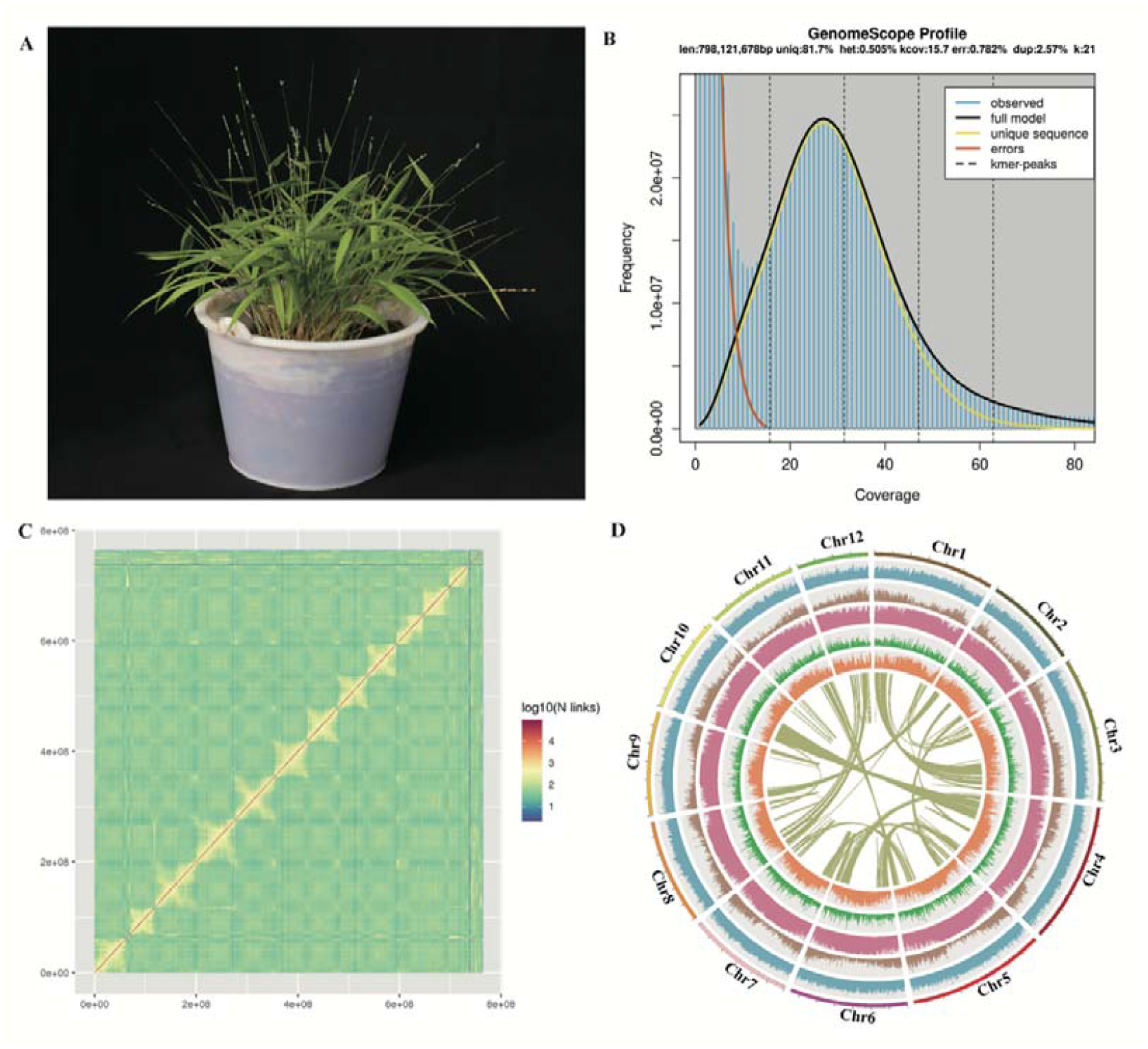
Summary of genome assembly and plant features of *O. granulata*. (**a**) Plant photo of *O. granulata*. (**b**) The K-mer (k=21) distribution and genome size estimation of *O. granulata* genome. (**c**) The genome-wide chromosomal interactions heatmap based on Hi-C data. (**d**) Circos ideogram of the *O. granulata* genome. Tracks from outer to inner include 12 pseudo-chromosomes, GC content, gene density, repeat element density, Copia-like element, and Gypsy-like element distribution. The innermost track indicates genomic synteny among the chromosomes.

**Fig 2.**
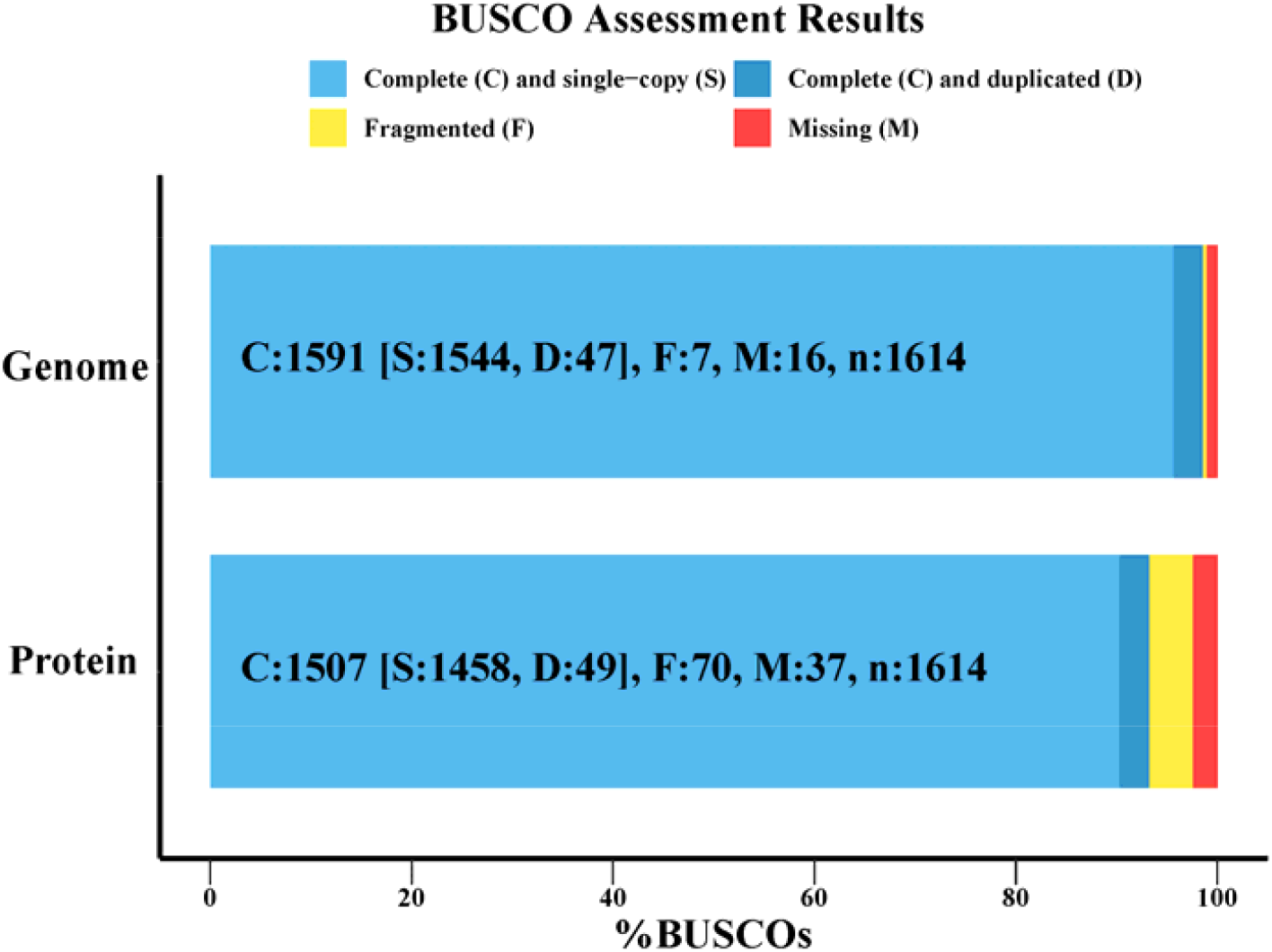
BUSCO completeness assessment of *O. granulata* genome assemblies and annotated protein-coding genes (PCGs).

**Fig 3.**
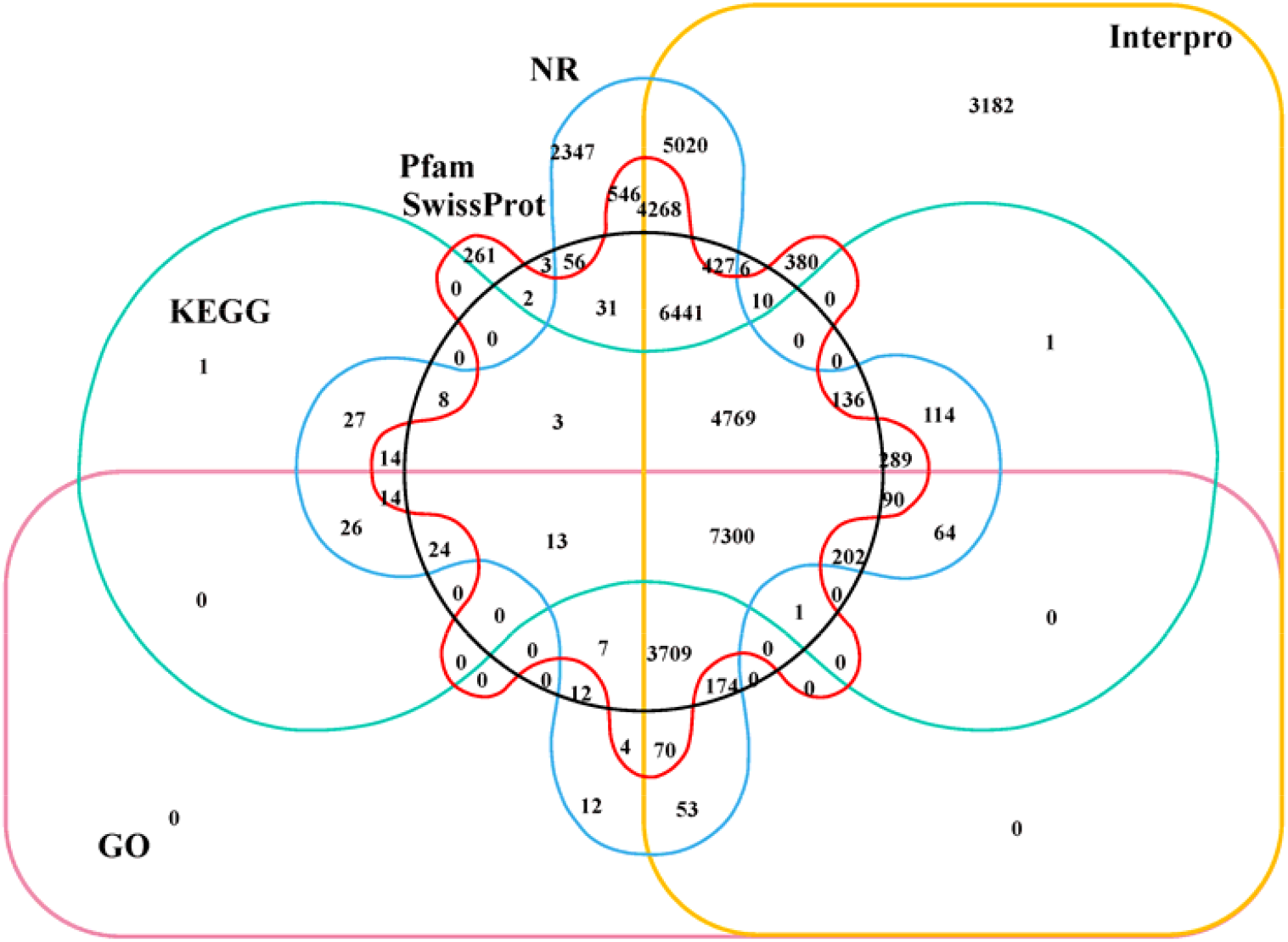
Venn diagram of gene functional annotations across multiple databases.

## Methods

### Plant materials, sample collection, and sequencing

For genomic DNA extraction, young healthy leaves of *O. granulata* were collected from Baoting City, Hainan Province, China, which were planted in the greenhouse of Hainan University, Haikou, China (**Fig. 1A**). Sampled leaves were immediately flash-frozen in liquid nitrogen and stored at -80 °C until further use. High molecular weight genomic DNAs (gDNAs) were extracted from leaves using improved CTAB method^25^ and evaluated using NanoDrop One spectrophotometer (NanoDrop Technologies, Wilmington, DE) and Qubit 3.0 Fluorometer (Life Technologies, Carlsbad, CA, USA). For the genome survey, the paired-end (PE 150bp) library was generated using the Illumina TruSeq DNA Nano Preparation Kit (Illumina, San Diego, CA, USA), and the library was sequenced on an Illumina HiSeq 2500 platform following the manufacturer’s instructions. As a result of Illumina sequencing, we obtained ∼103.03Gb of Illumina paired-end reads (**Table S1**). We performed a whole-genome shotgun sequencing (WGS) analysis with the single-molecule sequencing platform. This generated clean sequence data sets of ∼86.76 Gb with average read length of ∼11.58 Kb and yielded ∼113.56× coverage (**Table S1**). For Hi-C sequencing, formaldehyde was used for crosslinking the fresh leaves, and the crosslinking reaction was terminated using glycine solution. Subsequently, the Hi-C library was constructed based on the instructions and sequenced on the Illumina platform (Annoroad Gene Technology Co., Ltd), and ∼73 Gb raw reads were generated, which covered about 95.55× of the *O. granulata* genome (**Table S1**). The flag leaves, stem, panicles at the booting stage, panicles when flowering, panicles at the grain-filling stage, shoots of seedlings and roots of seedlings were collected for transcriptome sequencing. These tissue samples were rinsed using ddH_2_O and stored at −80°C until use after snap-freeze using liquid nitrogen with three biological replicates. Total RNA extraction was performed using the RNeasy Plant Mini Kit (Qiagen, Hilden, Germany). A cDNA library was built following the instructions, followed by paired-end sequencing on the NovaSeq platform (Illumina). A total of ∼35 Gb RNA-seq reads were obtained to assist the subsequent analysis of the *O. granulata* genome (**Table S1**).

### Chromosome-level genome assembly

Genome size of *O. granulata* was estimated from illumina data using k-mer frequency analysis. Jellyfish v2.3.032^26^ was first applied to extracting and counting canonical k-mer at k=21. Subsequently, GenomeScope^27^ was used to estimate the genome size from k-mer count data with parameters of “-k=21”. As a result, we estimated the genome size of *O. granulata* to be ∼798.12 Mb (**Fig. 1B**). The PacBio reads were *de novo* assembled by using FALCON v0.3.0^28^ with the following parameters: genome_size = 720000000, seed_coverage = 30, length_cutoff_pr = 5000, max_diff = 100, max_cov = 100. This consisted of six steps involving (1) raw reads overlapping; (2) pre-assembly and error correction; (3) overlapping detection of the error-corrected reads; (4) overlap filtering; (5) constructing graph; and (6) constructing contig. These processes produced the initial contigs. The assembly was then phased using FALCON-Unzip^28^ with default parameters. The primary contigs were polished as follows: firstly, quiver in SMRT Analysis version 2.3.0^29^ was used for genome polishing using PacBio data with a minimum subread length = 3000 bp, minimum polymerase read quality = 0.8. Paired-end sequencing reads were processed to remove adaptor and low-quality sequences using Trimmomatic v0.39^30^. Next, Illumina clean data from short libraries (≤ 500 bp) were aligned to the polished assembly using BWA v0.7.17^31^ with default parameters, and then, Pilon v1.24^32^ was used for sequence assembly refinement based upon these alignments. The parameters for pilon were modified as followed: –flank 7, –K 49, and –mindepth 15. Finally, we generated a primary assembly with a total length of ∼764.24 Mb, which spanned 95.74% of the genome size estimated by *k*-mer analysis (**Fig. 1C**; **Table 1**). The assembly comprised of 1,502 contigs with an N50 size of ∼1.22 Mb (**Table S2**). The cleaned Hi-C reads were mapped to the corresponding contigs using Juicer v1.9.9^33^. The unique mapped reads were taken as input for 3D-DNA pipeline v180114^34^ with parameters “-r 0”and then sorted and corrected manually using JuicerBox v1.11.08^35^. Manual review and refinement were performed to remove the potential errors. The gaps distributed among the pseudo-chromosome were filled with the PacBio raw reads using PBJelly2^36^ with parameter settings “-minMatch 8 -minPctIdentity 70 -bestn 1 - nCandidates 20 -maxScore −500 -nproc 10 –noSplitSubreads”. The assembly was subject to two rounds of Pilon v1.24^32^ polishing to remove the sequencing errors. The twelve pseudochromosomes were identified by distinct interaction signals in the Hi-C interaction heatmap (**Fig. 1C**), and the final assembled genome length was ∼737.24 Mb (**Fig. 1D**; **Table S3**), with a scaffold N50 of ∼59.32Mb, accounting for ∼96.47% of the assembled contigs (**Table 1**; **Table S3**). Compared to the ten other genome assemblies publicly available in the genus *Oryza*, the chromosome-level genome assembly of *O. granulata* obtained in this study showed remarkable sequence continuity and genome completeness (**Table S7**).

### Genome annotation and functional prediction

The repetitive elements in the *O. granulata* genome were identified by combining *de novo* and homology-based approaches. Tandem repeat sequences were annotated using Tandem Repeat Finder v4.09^37^ with default parameters. A total of six types (mono-to hexa-nucleotides) of simple sequence repeats (SSRs) were identified using the MISA v2.1^38^ identification tool with default parameters. For *de novo*-based searches, RepeatModeler v2.0.2a^39^, LTR_FINDER v1.07^40^, LTRharvest v1.5.9^41^, and LTR_retriever v2.9.1^42^ were applied for constructing *de novo* repeat libraries, by which RepeatMasker v4.1.3^43^ was employed to detect repeat sequences. For homology-based searches, we employed RepeatMasker v4.1.3^43^ against a known repeat library, Repbase v.19.06^44^. As a result, a total of ∼561.27Mb of repetitive elements occupying ∼73.46% of the *O. granulata* genome were annotated (**Table S5**). Most of these repeats were long terminal repeat (LTR) retrotransposons (∼56.20%) of the genome (**Table S5**). The DNA, LINE, and SINE classes accounted for ∼15.12%, ∼0.50%, and ∼0.04% of the genome, respectively (**Table S5**). Additionally, tRNAscan-SE v2.0^45^ software was used to predict tRNA genes. The rRNA, miRNA, and snRNA were predicted using INFERNAL v1.1.2^46^ software through searches against the Rfam database v15.0^47^. Finally, we annotated 651 miRNAs, 937 rRNAs, 994 tRNAs, and 608 snRNAs in the *O. granulata* genome (**Table 1**).

To annotate protein-coding genes in the *O. granulata* genome, gene models were obtained by combining the three approaches of *ab initio* gene predictions, homology-based predictions, and transcriptome-based predictions. The *ab initio* prediction was performed by AUGUSTUS v3.3.2^48^, SNAP v2013-11-29^49^, GeneMark-ES/ET^50^, GlimmerHMM v3.02^51^. For homology-based prediction, the Exonerate v2.2.0^52^ program was used to search against the protein sequences of *O. sativa* v7.0^53^, *Zea mays* RefGen V4^53^, *Arabidopsis thaliana* Araport11^53^, *O. longistaminata*^19^, *O. sativa* ssp. *japonica* cv. Nipponbare^54^, *O. rufipogon*^18^, *O. sativa* ssp. *indica* cv. HuaZhan^55^, and *O. sativa* ssp. *indica* cv. MingHui63^16^ genomes. For transcriptome-based prediction, Trinity v2.15.1^56^ was used for assembling transcripts based on RNA-seq data, and PASA v2.5.2^57^ software was employed for gene structure prediction based on transcriptome assemblies. Additionally, HISAT2 v2.2.1^58^ was employed for RNA-seq reads mapping onto the genome, and StringTie v2.2.1^59^ was used for the generation of transcript structure. The assembled transcripts were subsequently used for ORF (open reading frame) prediction using TransDecoder v5.5.0 (Haas, BJ. https://github.com/TransDecoder/TransDecoder). All predicted gene structures were integrated into a consensus set with EVidenceModeler v2.0.0^60^. Finally, 42,064 gene models were predicted after integrating the results of the three aforementioned methods (**Table 1**).

For the functional annotation of protein-coding genes, we aligned the predicted protein-coding gene sequences against public functional databases using BLAST v2.11.0^61^ (e-value < 1e-5), including Swiss-Prot^62^, NR^63^, KEGG, and KOG^64^. Gene Ontology (GO) was performed using InterProScan v5.55-88.0^65,66^. As a result, a total of 40,126 protein-coding genes were annotated for *O. granulata*, accounting for ∼95.39% of all predicted genes (**Fig. 3**; **Table S6**).

## Data Records

The Illumina short reads, PacBio SMART long-reads, Hi-C reads, genome assembly and annotation data were deposited in the National Genomics Data Center (NGDC)^67^, Beijing Institute of Genomics, the Chinese Academy of Sciences/China National Center for Bioinformation with BioProject accession numbers PRJCA061980^68^. The genome sequencing data were deposited in the Genome Sequence Archive (GSA) of NGDC under Accession Numbers CRA041662^69^. The genome assembly and annotation data were deposited in Genome Assembly Sequences and Annotations (GWH) of NGDC under accession number GWHISVE00000000.1^70^.

## Technical Validation

### Assessment of the genome assembly

The completeness of the assembled genome was evaluated using BWA v0.7.17^71^ and Benchmarking Universal Single-Copy Orthologs (BUSCO, v5.4.4) ^72^ with the embryophyta_odb10 lineage dataset. Approximately, ∼99.73% of the Illumina short reads were aligned to the genome, of which ∼94.10% of reads were properly mapped. The BUSCO analysis showed that the assembled genome sequences contained 1,591 (∼98.6%) complete BUSCOs, including 1,544 (∼95.7%) single-copy BUSCOs, 47 (∼2.9%) duplicated BUSCOs, and 7 (∼0.4%) fragmented BUSCOs (**Table S4**).

### Evaluation of the gene annotation

The annotated and integrated proteins were also evaluated using BUSCO v5.4.4^72^ with the lineage dataset embryophyte_odb10. Briefly, the proportion of complete core gene coverage was ∼93.3% (including ∼90.3% single-copy genes and ∼3.0% duplicated genes), and there were only a few fragmented (∼4.3%) and missing (∼2.4%) genes (**Table S4**), indicating high-quality annotation of the predicted gene models.

## Code availability

All software and pipelines were executed according to the manual and protocols of the published bioinformatic tools. All software used in this work is publicly available, with versions and parameters clearly described in Methods. If no detailed parameters were mentioned for a software, the default parameters suggested by the developer were used. No custom code was used during this study for the curation and/or validation of the datasets.

## Acknowledgements

We would appreciate Jia-jia Wan and Ju-jin Jia to grow the material and take the photo.

## Author contributions

Li-zhi Gao conceived and designed the study. Li-zhi Gao and Fen Zhang drafted the manuscript. Li-zhi Gao and Yue-hong Yang revised the manuscript. Yue-hong Yang collected samples. Fen Zhang and Wei Li executed data analysis. All authors read, edited, and approved the final manuscript.

